# CCLA: an accurate method and web server for cancer cell line authentication using gene expression profiles

**DOI:** 10.1101/858456

**Authors:** Qiong Zhang, Mei Luo, Chun-Jie Liu, An-Yuan Guo

## Abstract

Cancer cell lines (CCLs) as important model systems play critical roles in cancer researches. The misidentification and contamination of CCLs are serious problems, leading to unreliable results and waste of resources. Current methods for CCL authentication are mainly based on the CCL-specific genetic polymorphisms, whereas no method is available for CCL authentication using gene expression profiles. Here, we developed a novel method and homonymic web server (CCLA, Cancer Cell Line Authentication, http://bioinfo.life.hust.edu.cn/web/CCLA/) to authenticate 1,291 human CCLs of 28 tissues using gene expression profiles. CCLA curated CCL-specific gene signatures and employed machine learning methods to measure overall similarities and distances between the query sample and each reference CCL. CCLA showed an excellent speed advantage and high accuracy with a top 1 accuracy of 96.58% or 92.15% (top 3 accuracy of 100% or 95.11%) for microarray or RNA-Seq validation data (719 samples, 461 CCLs), respectively. To the best of our knowledge, CCLA is the first approach to authenticate CCLs based on gene expression. Users can freely and conveniently authenticate CCLs using gene expression profiles or NCBI GEO accession on CCLA website.

## Introduction

Cancer cell lines (CCLs) as important components offering unlimited biological materials play vital roles in life science studies. CCLs could serve as excellent model systems for the investigation of cancer biology, the simulation of drug response, and the development of clinical treatment on cancers (Holen et al. 2017). The utilization of CCLs is an effective and common practice in cancer researches (Bairoch A 2018; Barretina et al. 2012). However, the misidentification and contamination of CCLs are long-standing and prevalent problems (Capes-Davis and Neve 2016; Horbach and Halffman 2017; Development Organization Workgroup Asn-0002 2010), which could introduce erroneous, misleading, and false positive findings, and further result in invalid results and waste of resources. Researchers have raised extensive awareness of CCL authentication, the NIH and various journals have required cell line authentication for publications (Lorsch et al. 2014; Fusenig et al. 2017; Geraghty et al. 2014).

Up to date, available methods for CCL authentication were based on the DNA polymorphism information, such as short tandem repeats (STRs) and single nucleotide polymorphisms (SNPs) profiling (Dirks and Drexler 2005; Demichelis et al. 2008). STR profiling is the most common and standard method recommended by American Type Culture Collection (ATCC) for cell line authentication (Capes-Davis et al. 2010), and the SNP genotyping, either in combination with STRs or alone, was considered as an alternative method (Yu et al. 2015; Freedman et al. 2015). Although the STR and SNP methods had been widely used to authenticate CCLs in the past decades, they need additional experiments and could not be directly applied on expression data. Even though several methods (*e.g.*, CeL-ID) utilized RNA-Seq data to authenticate CCLs (Fasterius et al. 2017; Mohammad et al. 2019; Strong et al. 2014), their core algorithms still retrieved CCL-specific DNA polymorphism from RNA-Seq reads, which barricaded the application on gene expression data and required professional bioinformatics skills (*e.g*., SNP calling, polymorphism matching, and threshold). Thus, a convenient and precise tool using gene expression profiles for CCL authentication is an urgent requirement and will benefit the scientific reproducibility.

CCLs with similar genomic information have various expression profiles, which results in distinct characteristics for different CCLs (Domcke et al. 2013). The specifically expressed genes (SEGs), which were expressed in a unique or a small number of conditions, could serve as molecular features for different CCLs(Goodspeed et al. 2016; Zhang et al. 2018), and provide important clues for the CCL authentication. High-throughput transcriptome technologies including RNA-Seq and microarray have offered numerous expression data of CCLs, such as the Genomics of Drug Sensitivity in Cancer (GDSC) (Garnett et al. 2012), Cancer Cell Line Encyclopedia (CCLE) (Ghandi et al. 2019), Harmonizome (Rouillard et al. 2016), and others etc. (Klijn et al. 2015a; Hollingshead et al. 2014). These data provided convenience for the SEGs and marker genes detection in CCLs, and laid the foundation to develop methods for CCL authentication using gene expression profiles. Moreover, gene expression profiles based CCL authentication methods could bypass the procedure of DNA polymorphism calling, and benefit the authentication of CCLs which lack DNA information (*e.g.* transcriptional studies, gene function analysis, microarray data, and difficult to re-access the original cell lines etc.).

In this study, we developed a novel method and web server named CCLA (Cancer Cell Line Authentication), which combined machine learning methods and single sample gene-set enrichment analysis (ssGSEA) algorithm to authenticate 1,291 CCLs using gene expression profiles from RNA-Seq or microarray platform. Our evaluation results demonstrated that CCLA could rapidly and precisely authenticate CCLs.

## Results

### The summary of CCLA method

The workflow of CCLA is represented in the Figure 1 and the detailed algorithm is illustrated in the method section. In brief, CCLA integrated gene expression profiles and machine learning algorithms to authenticate the potential belonging for CCLs (Figure 1): 1) ssGSEA scores of signature gene sets were used as signatures for CCLs to replace the raw gene expression profiles, which could show a more robust pattern and avoid the severe bias of expression profiles from different sources; 2) A prediction model built by random forest (RF) algorithm was employed to pre-classify the query sample into a candidate category based on ssGSEA scores of signature genes; 3) After the categorization procedure, CCLA calculated the overall similarities and distances between the query sample and each reference CCL in the candidate category. Finally, the top 5 reference CCLs with the highest correlations and the least distances were considered as the potential belongings for the query sample.

**Figure 1.**
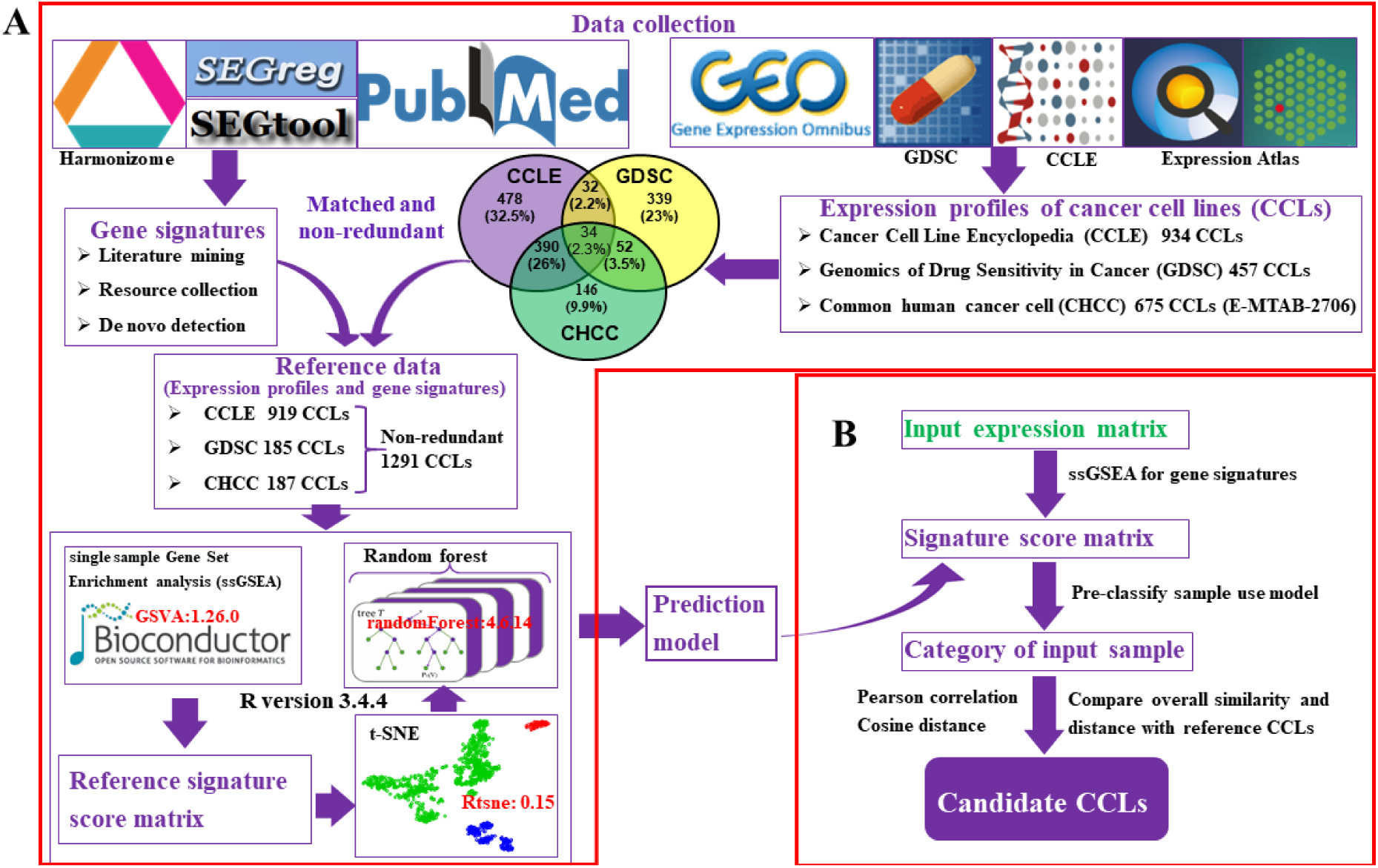
The data resources and algorithm of CCLA. (A) The resource collection and model construction for reference CCLs. The reference data of CCLs in CCLA were from three resources: CCLE, GDSC and CHCC (E-MTAB-2706 dataset in EBI). The gene signatures of CCLs were from three parts: 1) Text mining from publications; 2) SEGs collection from databases; 3) *De novo* detection by R package SEGtool. (B) The core steps of CCLA algorithm.

### Accuracy and feasibility assessment for CCLA on public datasets

To evaluate the performance of CCLA on CCL authentication, we used other datasets as test data which were independent of the reference one. We applied CCLA on three kinds of gene expression datasets from RNA-Seq and microarray platforms (Table 1), including: 1) Public untreated CCLs from different laboratories; 2) Different passages and treatments of CCLs; 3) Well-known or published incorrect and misidentified CCLs. In total, 719 samples of 461 CCLs from 15 individual studies were enrolled in this evaluation, including 573 samples of 456 CCLs from RNA-Seq technology and 146 samples of 14 CCLs from microarray platform (Table 1 and Supplementary Table S1). Among them, 511 samples were from GDSC database or E-MTAB-2706 dataset, which were shared by more than one sources. For example, the expression data of CCL “HCT15” were deposited in three databases, and the expression data in the CCLE database would be used as the reference profile, while the records in other two databases were worked as test data to assess the performance of CCLA. The confidence of CCLA results was mainly evaluated by the distributions of expressed signature genes in the query sample and resulting reference CCLs: 1) The profiles of expressed signature genes in the query sample and reference CCLs (Figure 2A, 2B); 2) The distribution of gene signatures in the query sample and the resulting reference CCLs (Figure 2C).

**Table 1.** Validation datasets and corresponding accuracies of CCLA.

| Datasets | #samples | #CCLs | Accuracy (%) |  |  | Exp. | Treatment |
| --- | --- | --- | --- | --- | --- | --- | --- |
|  |  |  | Top1 | Top3 | Top5 |  |  |
| E-MTAB-2706 | 399 | 399 | 93.98 | 96.49 | 97.24 | TPM | Untreated |
| GDSC | 112 | 112 | 81.25 | 87.50 | 93.75 | TPM | Drug |
| GSE32323 | 10 | 5 | 90.00 | 100 | 100 | Array | Drug and control |
| GSE54979 | 9 | 1 | 100 | 100 | 100 | Array | Drug and control |
| GSE55624 | 18 | 1 | 100 | 100 | 100 | Array | Drug and control |
| GSE66837 | 12 | 1 | 100 | 100 | 100 | Array | Drug and control |
| GSE73318 | 36 | 6 | 100 | 100 | 100 | RPKM | Drug and control |
| GSE83654 | 48 | 3 | 93.75 | 100 | 100 | Array | Drug and control |
| GSE101966 | 8 | 1 | 100 | 100 | 100 | FPKM | Knock out and control |
| GSE111485 | 18 | 1 | 100 | 100 | 100 | RPKM | Passages and control |
| GSE57820 | 12 | 1 | 100 | 100 | 100 | Array | miR-135b overexpression |
| GSE61692 | 4 | 1 | 100 | 100 | 100 | Array | Overexpression and control |
| GSE23655 | 12 | 1 | 100 | 100 | 100 | Array | Overexpression and control |
| GSE65168 | 8 | 1 | 100 | 100 | 100 | Array | Hypoxia and control |
| GSE7458 | 13 | 1 | 92.31 | 100 | 100 | Array | Degree and control |

**#samples:** The number of samples in the dataset; **#CCLs:** The number of CCLs in the study; **TopX:** The ratio of correct CCLs in the top X records in the outcome of CCLA; **Exp.:** The normalization method of gene expression data used in the study; **Treatment:** The treatment(s) used in the study.

**Figure 2.**
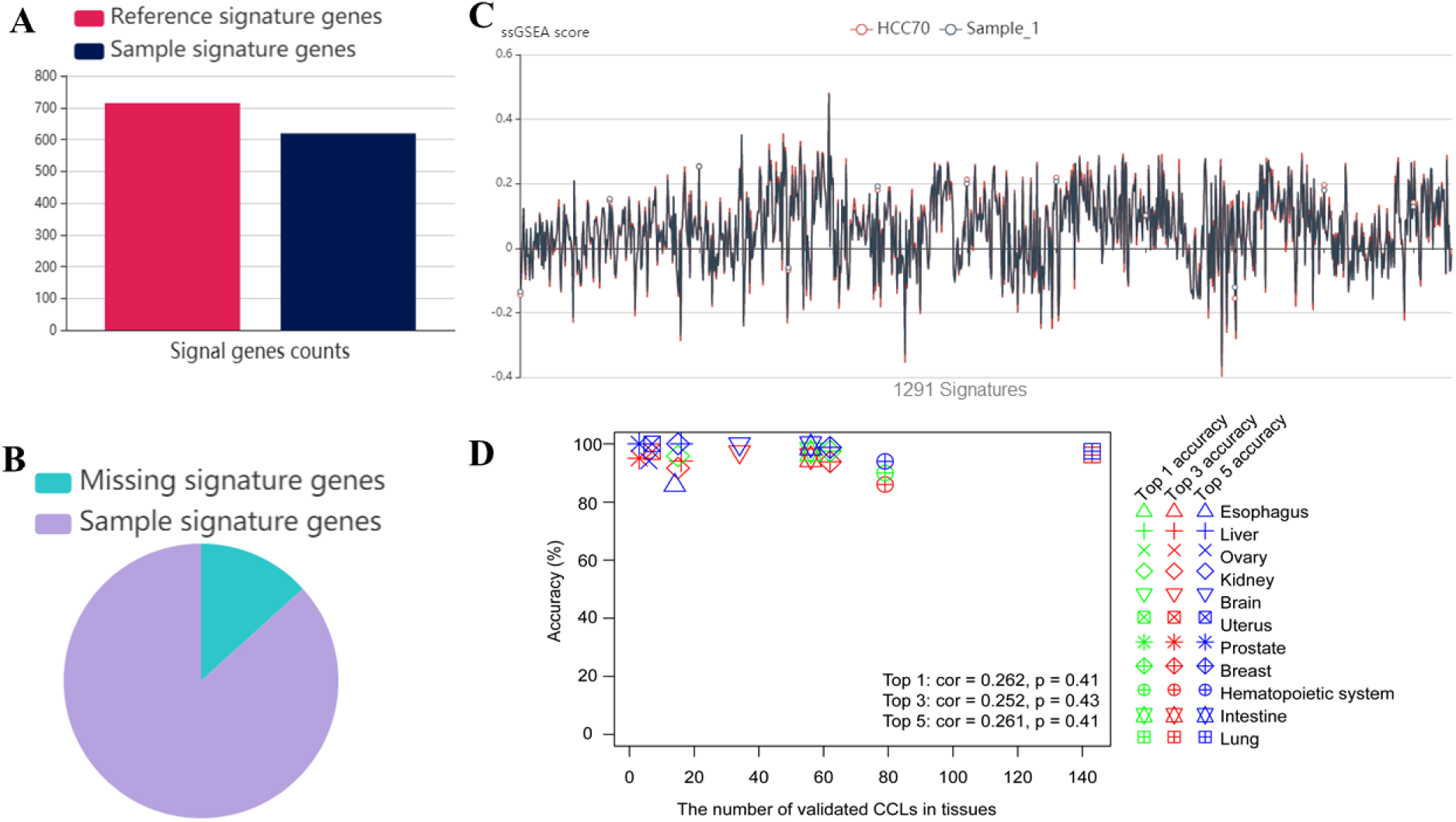
The accuracy assessment of CCLA. (A) The number of expressed signature genes in query sample and matched reference CCL; (B) The amount of missing signature genes in a query sample compared with the reference CCL; (C) The ssGSEA scores of signature gene sets in query sample and reference CCL. The X-axis indicates signatures of the reference CCL, while the Y-axis shows the ssGSEA scores of signature gene sets in the query sample (grey color) and candidate CCL (red color); (D) The accuracy of CCLA in different tissues. The Y-axis is the accuracy of validation datasets in corresponding tissue, and the X-axis shows the number of validated CCLs in the tissues. The top 1 accuracy means the target CCL ranks first of the outcomes of CCLA, while the top 3(5) accuracy indicates the target CCL appears in the first 3(5) results. The “cor” means the Pearson correlation coefficient between the accuracy and the reference CCLs in tissues, where the p is the P-value of the correlation. The low correlation here implies the accuracy of CCLA has no dependency with the number of reference CCLs per tissue.

As expected, CCLA showed a remarkable authentication power on CCLs both in the RNA-Seq and microarray datasets. Generally, CCLA achieved a high accuracy of 96.58% or 92.15% for the top 1 CCL (target CCL ranking the first one) on microarray or RNA-Seq data, respectively, while considering results in the top 3 list, the accuracy of CCL authentication was increased to 100% or 95.11% (Table 1 and Supplementary Table S1). The validation datasets for CCLA evaluation were widely spread in approximate 100 cancer types, suggesting that the power of CCLA was not limited in a small number of conditions.

Moreover, we wondered whether the number of reference CCLs per tissue could affect the authentication power of CCLA, and then investigated the relationship between the accuracy of CCLA and the number of reference CCLs in tissues (Figure 2D, Supplementary Table S2). To avoid the bias caused by the sample size of validation datasets, tissues (organs) with validation sample size more than 10 were enrolled in this evaluation. Notably, CCLA showed excellent performances (considering the top1, top3, top5 accuracy, respectively) on tissues containing different numbers (from 11 to 143) of reference CCLs (Figure 2D). The accuracy of CCLA showed a slight difference between tissues (no statistical significance) and did not increased (or decreased) with the number of reference CCLs in tissues (Figure 2D), suggesting there is no correlation between the number of reference CCLs per tissue and the accuracy of CCLA (Figure 2D, Pearson correlation coefficient < 0.27, P-value > 0.4).

Furthermore, we assessed the performance of CCLA on CCLs under different passages and treatments. The RNA-Seq dataset GSE111485 from GEO database containing 18 HeLa samples of different conditions [controls (n = 12), 7 passages (n = 3) and 50 passages (n = 3)] from different laboratories was employed to evaluate the authentication power of CCLA on CCLs with different passages. Although the passage times could influence the stability of genome and transcriptional profiles for CCLs(Liu et al. 2019), CCLA still showed a robust power on CCLs from different passages and laboratories. All of the 18 HeLa samples, no matter where they from and how many passages, were accurately authenticated as HeLa-original lines by CCLA (Supplementary Table S1). In addition, CCLA can perform well on expression data of CCLs under different treatments including drug treatment, gene over-expression, and microRNA transfection treatments etc. (Table 1). For example, 133 samples of CCLs treated by drugs from 6 independent studies were accurately authenticated as the original ones by CCLA (100% accuracy for top 3 results, Table 1), while the accuracy was slightly decreased in the samples from GDSC database (87.50% accuracy for 122 CCLs with drug treatment). Besides, CCLs with gene over-expression (GSE61692 and GSE23655) or gene knockout (GSE101966) treatments were all correctly authenticated as the original ones by CCLA (Table 1, Supplementary Table S1).

Furthermore, we also assessed the power of CCLA on the well-known misidentified CCLs, such as the MDA-MB-435 cell line, which was not a human breast cancer cell line but had been proved as M14 melanoma cell line by ATCC and several laboratories (Christgen and Lehmann 2007; Lacroix 2009; Prasad and Gopalan 2015). Interesting, the authentication for 8 MDA-MB-435 cell line samples (GSE128624) by CCLA showed that all of them were melanoma cell lines (Supplementary Table S3), implying the misidentification of MD-AMB-435 cell line was a long-time event and CCLA could serve as a valuable tool to benefit the reproducibility of scientific data and results based on the available expression data.

### Comparison with other approaches

Although a few methods (*e.g.*, CeL-ID and Fasterius’ method) could utilize RNA-Seq data to authenticate CCLs (Fasterius et al. 2017; Mohammad et al. 2019), their core algorithms retrieved genomic polymorphism of samples from RNA-Seq reads (not the expression profiles) to match CCL-specific SNPs and could not be applied on microarray data. Meanwhile, these methods just stated a pipeline and did not provide any mature software (package, tool or online server) and important parameters (e.g. the version of used tools, the match pattern, the reference SNPs of CCLs, and the threshold etc.) in their publications, which made it very difficult to reproduce their results. Thus, we just compared the authentication results of CCLA and CeL-ID based on the same RNA-seq data used by CeL-ID (Table 2).

**Table 2.** The comparison of CCLA with CeL-ID.

|  | CCLA | CeL-ID |
| --- | --- | --- |
| Data type | Expression data | SNPs from RNA-Seq reads |
| Support platform | RNA-Seq & microarray | RNA-Seq |
| Expression data | Y | N |
| Time | 10s | Excessive time |
| Pre-condition | None | Professional skills and several software<br>for SNP calling and pattern match |
| Convenience | Web server | No mature tool |
| Capacity | 1,291 | unknown |
| Cost | Free | Free |

Two datasets containing 20 samples (12 samples of MCF7 CCL from GSE23655, 8 samples of HCT116 CCL from GSE101966) were enrolled to benchmark the performance of CCLA and CeL-ID. We first processed the RNA-Seq data to obtain gene expression profiles of CCLs according to the HISAT2-StringTie protocol (Pertea et al. 2016), and then applied CCLA to authenticate them. All samples in GSE101966 dataset were authenticated as HCT116-orignal cell lines, while samples of MCF7 cell line were authenticated as MCF7 as well. Thus, our benchmark results suggested that CCLA showed a similar accuracy as the method CeL-ID using DNA polymorphism from RNA-Seq data (Table 2). Moreover, CCLA showed several excellent advantages on time and convenience (time: a few seconds for CCLA, much time cost for SNP calling in CeL-ID; cost: free; precondition: only need gene expression profiles for CCLA, several bioinformatics tools need in CeL-ID; polymorphism loss: none for CCLA, always issues for polymorphism based methods; and etc.). Furthermore, our CCLA is the only available tool and online web server to provide mature and convenient service for CCL authentication using gene expression profiles.

### Website interface of CCLA

For the convenient application of CCLA by users, we developed a homonymic web server to provide free service of 1,291 CCLs authentication (Figure 3). Users could easily authenticate and assess their interested CCLs using gene expression data. CCLA accepts a NCBI GEO accession of microarray data or unfiltered gene expression matrix (Figure 3A), whose rows represent the normalized expression value for genes (FPKM, RPKM and TPM format for RNA-Seq data, while RMA and MAS5 for microarray data) and columns are samples. Once the target CCL is selected (Figure 3A), CCLA provides an overall view of outputs and evidence for the authentication of query samples (Figure 3B). For example, an individual page displays the detailed results: 1) The top five candidate CCLs for each query sample (Figure 3C); 2) The profiles of expressed signature genes in the query sample and reference CCLs (Figure 2A, 2B); 3) The gene signal distribution of the query sample and the resulting reference CCL to the query sample evaluated by Pearson correlation and cosine distance (Figure 2C); 4) The expression pattern of each signature gene in the query sample and the resulting reference CCL (Figure 3D). Our CCLA is freely available at http://bioinfo.life.hust.edu.cn/web/CCLA/.

**Figure 3.**
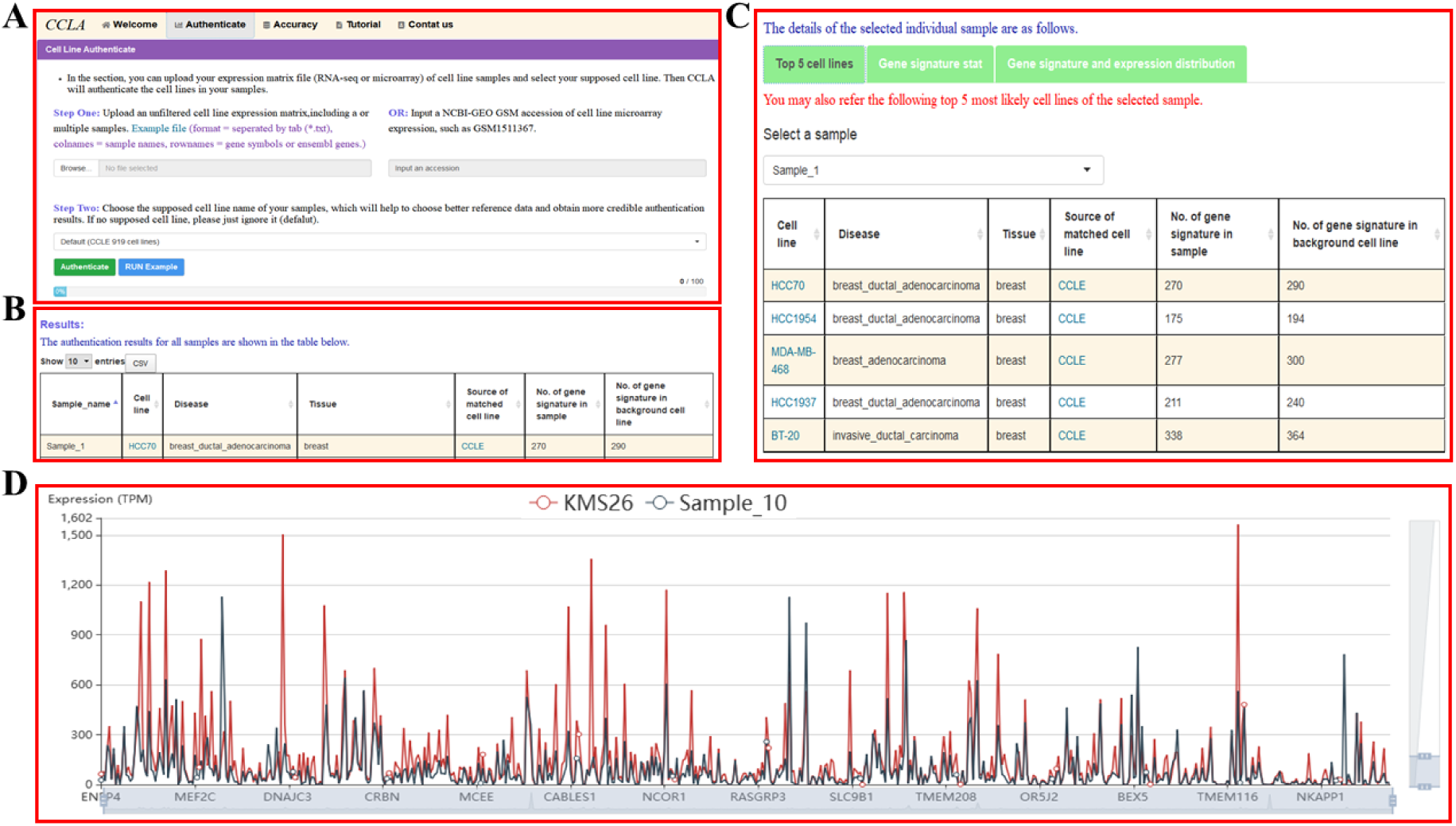
The interface of CCLA web server. (A) A snapshot of the authentication page; (B) A partial of the authentication result page for the query sample in CCLA web server; (C) The detail results of top 5 candidate CCLs for the query sample; (D) The expression profiles of signature genes in the query sample and the reference CCL.

## Discussion

CCLs derived from human cancers are important biomaterials for cancer biology exploration, pre-clinical modeling, clinical application and drug validation (Goodspeed et al. 2016; Wilding and Bodmer 2014). The misidentification and mislabeling of CCLs are long-standing and widespread problems in biomedical researches for decades (Vaughan et al. 2017; Christgen and Lehmann 2007; Jäger et al. 2013), and large-scale cross-contaminations and misidentification of CCLs were reported recently (Horbach and Halffman 2017; Strong et al. 2014; Teixeira da Silva 2018; Rebouissou et al. 2017; Bairoch 2018). However, available methods for CCL authentication were based on DNA polymorphism, which could not be well applied on the transcriptome datasets and be too cumbersome for biomedical researchers. To address these concerns, we developed CCLA using gene expression profiles to rapidly authenticate CCLs with high accuracy and robustness. Furthermore, we built a homonymic web server to provide free CCL authentication for researchers (http://bioinfo.life.hust.edu.cn/web/CCLA/).

The authentication of cell lines is a key factor for the reliability of biomedical researches, which is required for the grant application and manuscript publication (Lorsch et al. 2014; Potash and Anderson 2009). DNA polymorphism (*e.g.* STRs and SNPs) based approaches for CCL authentication analyze the similarities between the query sample and reference CCLs in specific loci. Even there are high-throughput sequencing data for CCLs, the loci-specific polymorphisms needed for CCL authentication were often omitted from sequencing or quality control procedures or uncertain RNA editing (Mohammad et al. 2019; Capes□Davis et al. 2013; Richards et al. 2015; Otto et al. 2017), and different sequencers or different variant calling pipelines may generate substantial disagreement results (Hwang et al. 2015, 2019; Coudray et al. 2018). Meanwhile, due to the severe genomic instability of CCLs and heterogeneous NGS profiles, the coincidence of genetic polymorphism from different laboratories and research projects was less than expected (Hudson et al. 2014; Alkan et al. 2011), thus different algorithms or workflow designs for the authentication of the same CCL were required. Moreover, the excess passages and environmental conditions (*e.g.* drug exposure) could lead to the acceleration of genetic drift and alteration of alleles information for CCLs, which may require special algorithms and interpretation for the profiles of STR or SNP from unstable CCLs (Eltonsy et al. 2012; Marx 2014) and pose another challenge for the authentication methods using DNA polymorphism (Eltonsy et al. 2012). Additionally, large number of CCLs used in previous studies were focused on the functional study of genes or pathways and the alterations of transcriptional profiles under specific conditions, which lacked enough genomic polymorphisms for CCL authentication (*e.g.* microarray and RNA-Seq data). Our CCLA implemented GSVA algorithm to calculate a robust signal score matrix for CCLs, and then employed machine learning approaches to further identify the belongings of input dataset. Instead of a fixed panel of limited number of STRs or SNPs, CCLA utilized a stable signal matrix of gene expression profiles to represent CCLs and avoid the bias caused by different passages of CCLs and the genome instability (Table 1), which in turn strengthen the authentication power on the experimental treated CCLs (Table 1).

CCLA achieved an excellent authentication power for CCLs both on the RNA-Seq and microarray data (Table 1). The results of CCLA from comprehensive validation data (719 samples of 461 CCLs in 21 tissues from 15 independent datasets) demonstrated that CCLA could authenticate CCLs with high precision: 92.15% (528/573), 95.11% (545/573) of top1, top3 accuracy for RNA-Seq data; 96.58% (141/146), 100% (146/146) of top1, top3 accuracy for microarray data (Table 1). Furthermore, CCLA performed well on CCLs with different passages or drugs or gene manipulation treatments (Table 1), suggesting the robustness of CCLA on expression data of various treatments. Additionally, our validation results showed that CCLA had a good sensitivity and accuracy on distinguishing CCLs from the same tissue origin (Figure 2D and Supplementary Table S2). In this way, CCLA is an essential tool to integrate metadata and ensure the reproducibility and reliability of results from cancer research using CCLs of previous studies.

Although CCLA showed a high accuracy for the authentication of 1,291 CCLs, the contamination (such as mixed with other cell lines and the *Mycoplasma*) remained a serious problem uncovered in this study. The issue of contamination with other cell lines often exists without obvious signs in experiments, and could result in global alteration of signal scores for the donor cell line. Considering the core algorithm, CCLA may not perform well on the cross-contamination conditions, while the DNA polymorphism based methods may be a better choice for this case. The contamination of *Mycoplasma* could influence cell metabolism and growth, induce chromosomal abnormalities, and alter transcriptome profiles (Geraghty et al. 2014; Olarerin-George and Hogenesch 2015). Our results demonstrated that ∼30% (21/60) CCLs with low-level *Mycoplasma* contamination were identified to their original ones, whereas nearly 70% (39 out of 60) CCLs with severe *Mycoplasma* contamination were authenticated as others (Supplementary Table S4). One possible reason is that the expression patterns of CCLs with severe *Mycoplasma* contamination were significantly changed, which was reported by previous studies (Olarerin-George and Hogenesch 2015; Zhang et al. 2006). In this manner, CCLs with severe contamination of *Mycoplasma* may be authenticated as different one by CCLA, and CCLA could serve as an indirect approach to imply the contamination of mycoplasma (or perhaps used the wrong CCL). Finally, CCLA consolidated 1,291 commonly used CCLs in this version and we will keep updating with the increase of standard datasets. No a single method could provide all of the information for human cell line authentication (Development Organization Workgroup Asn-0002 2010), and our CCLA could represent the valuable candidate to identify CCLs on gene expression data.

The authentication of CCLs is an essential issue to avoid fake data and ensure the scientific reproducibility and credibility. Although DNA polymorphism profiling based methods are recommended for CCL authentication (Development Organization Workgroup Asn-0002 2010), the cost and inconvenience of these methods and the physical re-access for the original CCLs appear as main roadblocks for their universal applications on CCLs of previous studies (Freedman et al. 2015). Our transcriptome profiles based method CCLA could be an important supplemental approach and new direction. Additionally, most of available methods were not user-friendly for researchers because they need extra bioinformatics and programming skills. Our CCLA offered a convenient web server for the scientific community to rapidly authenticate CCLs and valuable references for journals with less time, money and effort, and even shed new light for the transcriptome profiles based cell line authentication.

## Conclusion

In summary, CCLA is freely available and will largely contribute to the decrease of CCLs misidentification. To the best of our knowledge, CCLA is the first approach and the first online website to authenticate CCLs using gene expression data. CCLA can serve as a useful resource for cancer research and improve the reliability of biomedical results.

## Materials and Methods

### Collection for gene expression profiles of non-redundant reference cancer cell lines (CCLs)

To obtain the relatively unbiased and authoritative gene expression profiles of reference CCLs, we curated the RNA-Seq gene expression profiles of CCLs from 3 generally recognized CCL resources: 1) Cancer Cell Line Encyclopedia (CCLE), which contains expression profiles of 934 CCLs from RNA-Seq data(Ghandi et al. 2019); 2) Genomics of Drug Sensitivity in Cancer (GDSC), which deposits the expression profiles of 457 CCLs from RNA-Seq data(Yang et al. 2013); 3) The E-MTAB-2706 dataset, which is a comprehensive transcriptional portrait of 675 common human CCLs (Klijn et al. 2015b).

Furthermore, we examined the integrity of information for all the CCLs above. Briefly, all the introductions of reference CCLs were retrieved using an in-house “web crawler” script programmed by the python language and its libs (e.g. urllib, BeautifulSoup, and requests etc.). First, CCLs with a similar character string (e.g. “HCT 116” or “HCT116” or “HCT-116” or “HCT_116”, but not limited in this style) and the same origin (e.g. from the colon or large-intestine etc.) were deemed as the same kind of a CCL with different aliases. Then CCLs with similar origins but large-distance of their names (20%, e.g., the character difference between SW1417 and SW1463, not limited in this situation) were carefully checked and manually examined from the webpages of the resources. In addition, when a CCL was stored in more than one source, the priority of its gene expression profile as a reference in CCLA was ranked by the following order (CCLE > GDSC > E-MTAB-2706). For example, the CCLE and GDSC databases simultaneously collected the expression profiles of HCT116 CCL, and in this case, the gene expression profile of HCT116 CCL in the CCLE resource was served as a reference CCL, while the one in the GDSC was used as a validation sample for HCT116 CCL. Based on the above procedures, 1,471 kinds of non-redundant or unique reference CCLs (883 CCLs from CCLE database, 391 from GDSC, and 146 from E-MTAB-2706) were kept for further analyses.

### Curation of signature genes for CCLs

First, the gene signatures of each CCL were retrieved from literature mining, resource collection and de novo detection processes: 1) Literature mining from publications. In this process, we used several key words (e.g., “maker gene”, “specifically expressed gene (SEG)”, and “highly expressed gene” etc.) in the PubMed database to retrieve candidate signature genes for corresponding CCLs; 2) Resource collection. Two databases Harmonizome and SEGreg (Rouillard et al. 2016; Tang et al. 2018) were the main resources to collect the signature genes. In Harmonizome, those candidate signature genes with a score > 1 were used, which indicates that the gene has a strong positive gene-CCL association. In SEGreg database, genes with the tag “high” in the corresponding CCL was deemed as candidate signature genes; 3) De novo detection, SEGs were detected using SEGtool (Zhang et al. 2018) (default parameters, p-value <= 0.05, highly expressed pattern) on gene expression profiles of 1,471 reference CCLs, and the output SEGs were acted as candidate signature genes as well.

Second, candidate signature genes from the above three processes were integrated to explore putative signature genes by the following two steps: 1) For CCLs from the same tissue (or organ), we calculated and adjusted the ratio of tissue-specific genes to candidate signature genes. For example, if the ratios of tissue-specific genes (with the number of 30) were more than 40% in 5 CCLs, we randomly assigned the same number of tissue-specific genes (e.g., the number is 30/5 = 6 in this case, allowed 1/5 = 20% repetition) to the 5 CCLs; 2) For CCLs with similar candidate signature genes, we implemented the same operation as the step 1. Furthermore, we measured the reliability of signature genes in reference CCLs by examining their expression levels. After the above processes, the retained genes were considered as putative signature genes. Finally, 180 out of 1,471 CCLs that did not have enough signature genes (less than 50) were dropped, and the rest 1,291 reference CCLs were kept for further analyses.

### Signature calculation and model construction for CCLs

To avoid the bias and technical variability of gene expression caused by the noise, different quantile normalization methods, and various experiment treatments, ssGSEA algorithm was implemented to calculate the enrichment scores of signature genes for each CCL, which could serve as robust expression features compared with the raw gene expression profiles (Figure 1). Thus, the raw expression profiles of reference CCLs have been represented by ssGSEA scores of signature genes sets, and each CCL has the same 1,291 signatures, whose expression values are ssGSEA scores. Next, we constructed a 1,291 × 1,291 signature matrix for the reference CCLs, in which, each row is the corresponding signature values in 1,291 CCLs, and columns represent CCLs. Furthermore, t-distributed stochastic neighbor embedding (t-SNE) algorithm was used for the classification and clustering of reference CCLs based on their signatures (parameters: dims = 3, perplexity = 50, max_iter = 5000, theta = 0, pca = TRUE), and three groups were obtained. Subsequently, we employed the random forest (RF) algorithm to extract features from reference CCLs with their group labels determined by t-SNE (the importance of each feature was represented in the Supplementary Table S5), and then built a prediction model that would been applied to estimate the potential group for the query CCL (Figure 1).

### CCL authentication

In order to accurately authenticate CCLs, CCLA calculates the ssGSEA score of signature genes for the query CCL, then applies the prediction model (built by RF algorithm in the model construction step above) to pre-classify the candidate group of the query CCL (Figure 1). Then CCLA employs Pearson correlation and cosine distance to measure the similarities and divergences between the query CCL and each reference CCL in the pre-classified category. Then, CCLA ranked reference CCLs in the given category by Pearson correlation coefficient and cosine distance. The reference CCL with the highest similarity and least divergence was considered as the putative belonging of the query CCL, and the top 5 CCLs were also listed as candidate results.

### Validation data collection

Both gene expression profiles of CCLs from RNA-Seq and microarray platforms were curated to evaluate the accuracy of CCLA. Three kinds of CCLs with gene expression profiles were collected: 1) Public untreated CCLs from different laboratories; 2) Different passages and treatments of CCLs; 3) Well-known misidentified CCLs (Table 1).

We employed the following criteria to judge a successful authentication in CCLA: 1) The consistency between paper declared CCL and the results of CCLA. For example, if a CCL was identified as another one by CCLA which was different from the original paper, we deemed this as an inaccuracy authentication, otherwise is correct, expect for the well-known misidentified or contaminated CCLs (such as MDA-MB-435 cell line, the American Type Culture Collection (ATCC) reported that the MDA-MB-435 cell line is not breast cancer but actually melanoma related cell line); 2) For the well-known misidentified CCLs (e.g. MDA-MB-435 cell line), all the MDA-MB-435 strains were considered as the melanoma cell lines, and if any MDA-MB-435 cell line was identified as the melanoma origin, we deemed this authentication was a correct case.

## Abbreviations

CCLA: cancer cell line authentication
CCL: cancer cell line
GDSC: Genomics of Drug Sensitivity in Cancer
CCLE: Cancer Cell Line Encyclopedia
CHCC: common human cancer cell
EBI: European Bioinformatics Institute
FPKM: fragments per kilobase per million mapped fragments
RPKM: reads per kilobase per million mapped reads
TPM: transcripts per kilobase per million mapped reads
ssGSEA: single sample gene-set enrichment analysis
SEG: specifically expressed gene
SNP: single nucleotide polymorphism
STR: short tandem repeat

## Authors’ contributions

Q.Z and M.L: Methodology, Data collection, Webserver work, and Manuscript writing; M.L, Q.Z and C.J.L: bioinformatics analysis; AG and QZ: Conceptualization, Writing, Revising, Funding Acquisition, and Supervision.

## Acknowledgements

We would like to thank colleagues for data production and web server testing. We are also grateful to our users and all members of our lab for their valuable suggestions and comments.

We also acknowledge the research funding from the National Key Research and Development Program of China (Grant No. 2017YFA0700403), National Natural Science Foundation of China (NSFC) (Grant Nos. 31822030, 31801113, and 31771458), China Postdoctoral Science Foundation (Grant No. 2018M632830 and 2019M652623), the Fundamental Research Funds for the Central Universities (2018KFYRCPY002) and the program for HUST Academic Frontier Youth Team.

## Supporting Information

**Supplementary Table S1: Detailed results of authenticated CCLs for the test datasets.**

**Supplementary Table S2: The accuracy of CCLA on different tissues.**

**Supplementary Table S3: The authentication results of MD-AMB-435 datasets.**

**Supplementary Table S4: The authentication results of CCLs contaminated by *Mycoplasma.***

**Supplementary Table S5: The importance of features in the predicted model from RF algorithm.**

